# Hsp70 inhibits aggregation of Islet amyloid polypeptide by binding to the heterogeneous prenucleation oligomers

**DOI:** 10.1101/2020.03.30.016881

**Authors:** Neeraja Chilukoti, Bankanidhi Sahoo, S Deepa, Sreelakshmi Cherakara, Mithun Maddheshiya, Kanchan Garai

## Abstract

Molecular chaperone Hsp70 plays important roles in the pathology of amyloid diseases by inhibiting aberrant aggregation of proteins. However, mechanism of the interactions of Hsp70 with the amyloidogenic intrinsically disordered proteins (IDPs) is not clear. Here, we use Hsp70 from different organisms to show that it inhibits aggregation of Islet amyloid polypeptide (IAPP) at substoichiometric concentrations even in absence of ATP. The effect is found to be the strongest if Hsp70 is added in the beginning of aggregation but progressively less if added later, indicating role of Hsp70 in preventing primary nucleation possibly *via* interactions with the prefibrillar oligomers of IAPP. Fluorescence Correlation Spectroscopy (FCS) measurements of the solutions containing fluorescently labelled Hsp70 and IAPP exhibit fluorescence bursts suggesting formation of heterogeneous complexes of oligomeric IAPP binding to multiple molecules of Hsp70. Size exclusion chromatography and field flow fractionation are then used to fractionate the smaller complexes. Multiangle light scattering and FCS measurements suggest that these complexes comprise of monomers of Hsp70 and small oligomers of IAPP. However, concentration of the complexes is measured to be a few nanomolar amidst several μmolar of free Hsp70 and IAPP. Hence, our results indicate that Hsp70 interacts poorly with the monomers but strongly with oligomers of IAPP. This is likely a common feature of the interactions between the chaperones and the amyloidogenic IDPs. While strong interactions with the oligomers prevent aberrant aggregation, poor interaction with the monomers avert interference with the functions of the IDPs.

## Introduction

Pathology of type 2 diabetes mellitus (T2DM) is characterized by dysfunction of pancreatic β-cells, progressive decline in insulin secretion and loss of the β-cell mass. Aetiology of β-cell damage is multifactorial (1), however, growing evidences suggest that amyloid aggregation of the Islet amyloid polypeptide (IAPP) is a major cause of death of the β-cells, particularly at the late stage of T2DM (2,3). IAPP, a 37-residue peptide hormone, is co-expressed and co-secreted with insulin from the pancreatic β-cells. The triggers for aggregation of IAPP in the pancreas is not clear but several different mechanisms such as defects in IAPP biosynthesis, folding, trafficking, and/or processing in response to insulin resistance and inflammation have been proposed to be responsible (4,5). *In vitro*, IAPP aggregates into soluble oligomers and insoluble fibrils at micromolar concentrations (6). Aggregates of IAPP, particularly the soluble oligomers are considered to be the major toxic species affecting cellular integrity (7–9).

Heat shock protein Hsp70 is a central component of the cellular chaperone system playing essential roles in protein homeostasis under normal and stress conditions by preventing protein misfolding and aggregation (10,11), and protecting the cells from the cytotoxicity caused by amyloid aggregation (12). Expression of Hsp70 is found to be lower in T2DM in the insulin sensitive tissues (13,14). Furthermore, overexpression of Hsp70 in obese mice can delay or prevent the symptoms of diabetes such as inflammation and insulin resistance (15). Chaperone complementation studies either by overexpression or by exogenous addition are shown to minimise the cytotoxic effects of the amyloids in model organisms of T2DM and other amyloid diseases (16,17). Furthermore, Hsp70 is found co-deposited with IAPP in the islet amyloids in T2DM (18). Therefore, elevation of the levels of Hsp70 and also improving its specificity for IAPP are being proposed as therapeutic strategies in T2DM (16–19,20).

*In vitro*, Hsp70 along with the cochaperones Hsp40 and the nucleotide exchange factor (NEF) have been reported to inhibit aggregation of amyloid-β, α-synuclein, polyglutamine polypeptides, tau protein and IAPP (21–25). Furthermore, the Hsp70 machinery can also cause disaggregation of the soluble oligomers and/or the insoluble fibrils in presence of ATP (26). The inhibition of aggregation and disaggregation of preformed aggregates requires only substoichiometric concentrations of Hsp70. Hence, Hsp70 is hypothesized to act in an ATP dependent catalytic cycle, which involves binding to the exposed hydrophobic patches of the ‘misfolded’ or the ‘aggregation competent’ conformers of the globular proteins, refolding it, and then releasing it in the ‘non-amyloidogenic’ monomeric forms (27). Since intrinsically disordered proteins (IDP) do not possess a stable structure it is probably not meaningful to think about a ‘misfolded’ IDP. Hence, it is unclear how its interaction with the chaperones is regulated. Recent studies have shown ATP independent inhibition of amyloid aggregation by Hsp70 and several other chaperone proteins (28,29). For example, Loomes et al have reported that Hsp70 delays aggregation of IAPP at substoichiometric (1:200) concentrations even in absence of ATP (25). Therefore, inhibitory action of Hsp70 does not necessarily require the catalytic activity. However, substoichiometric nature of the interactions indicates that Hsp70 must interact with one or more on-pathway intermediates of IAPP. However, biophysical characterization of these intermediates has been challenging due to its transient and heterogeneous nature (30). Furthermore, Hsp70 is also known to exist as monomers, dimers and higher oligomers (31). Hence, little is known about the properties of the Hsp70/IAPP complexes. We think that investigations using single molecule techniques can help to overcome the limitations posed by the heterogeneity and the transient nature of the interacting species.

Here we have used three different Hsp70 chaperone proteins, such as, *E. coli*. DnaK (UniProt ID-P0A6Y8), *Mycobacterium tuberculosis* Hsp70 (MTB-Hsp70, UniProt ID-P9WMJ9) and human Hsc70 (UniProt ID-P11142) and examined its inhibitory effects on different phases, *viz*, the lag and the growth phase of the aggregation of human IAPP. Then we have measured the size of the Hsp70/IAPP complexes using Multiangle light scattering (MALS) and Fluorescence Correlation Spectroscopy (FCS). Size Exclusion Chromatography (SEC) and Field Flow Fractionation (FFF) are employed to ameliorate the heterogeneity of the complexes. Our results indicate that Hsp70 binds to the prenucleation oligomers to inhibit the growth of the amyloids of IAPP.

## Results

### Inhibition of amyloid aggregation of IAPP by Hsp70 proteins

First, we examined whether Hsp70 inhibits aggregation of IAPP in absence of ATP. Figure 1A-C show that time course of aggregation of IAPP (20 μM) is characterized by a lag (t = 0-4 h), a growth (t = 4-12 h) and a saturation phase (t >15 h) consistent with the nucleation dependent aggregation of amyloid proteins (32). Addition of substoichiometric concentrations (0, 0.125, 0.25 and 0.5 μM) of DnaK, MTB-Hsp70 or Hsc70 delays the kinetics of aggregation dramatically. While all the three chaperones inhibit amyloid aggregation, there are some differences. For example, the inhibitory effects appear to be the strongest in presence of DnaK and weakest in presence of Hsc70. The sequence identity between the Hsp70 proteins used here is about 50% (33). Therefore, despite large differences in the primary sequences, Hsp70 across different species share common mechanisms in protection against amyloid aggregation. This is consistent with the highly conserved nature of the overall structural organization of the variants of Hsp70 (33).

**Figure 1:**
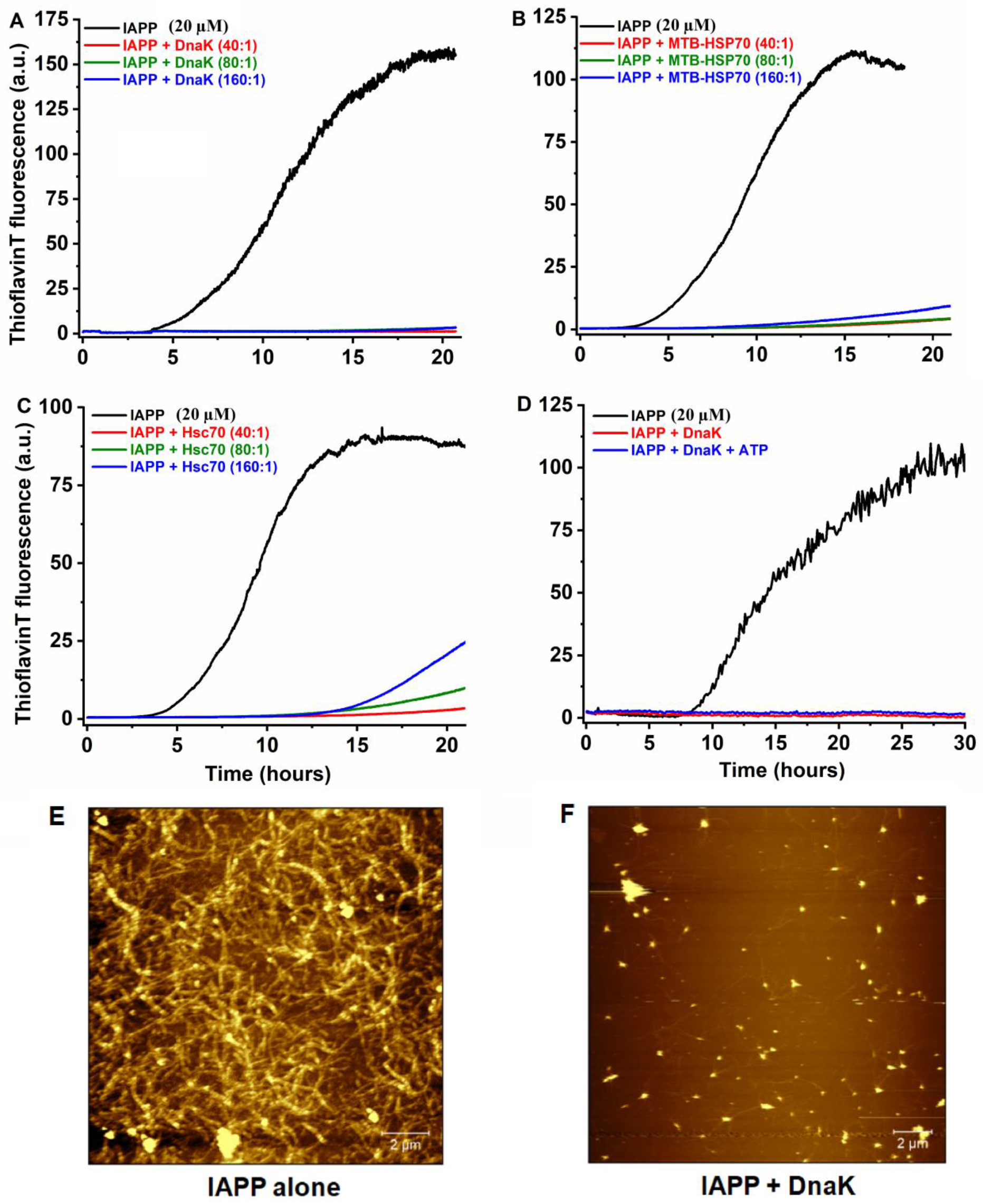
Effect of Hsp70 on aggregation of IAPP. A-C) Kinetics of aggregation of 20 μIAPP in presence of substoichiometric concentrations of *E.coli* DnaK (A), MTB-Hsp70 (B) and human Hsc70 (C). Ratios indicated in the figures are molar ratios. D) Effect of DnaK (0.5 μM) on the kinetics of aggregation of 20 μM IAPP in absence or presence of 1 mM ATP. Aggregation is monitored using fluorescence of Thioflavin T. E-F) AFM images of the 20 μM IAPP incubated for 48 hours in absence or presence of 0.5 μM DnaK. All experiments are performed in PBS buffer containing 1 mM EDTA and 1 mM sodium azide at pH 7.4 and incubated at 37 °C with continuous stirring.

Figure 1D examines whether the inhibitory effects of Hsp70 could be enhanced in presence of ATP. It may be seen that the inhibitory effects of DnaK on the kinetics of aggregation of IAPP in presence and absence of ATP are quite similar. This is not surprising because inherent ATPase activity of Hsp70 is known to be poor (11).

Some literature reports suggest that Hsp70 prevents amyloid formation, but it promotes amorphous aggregation (28). We have performed Atomic Force Microscopy (AFM) measurements on the aggregates of IAPP prepared upon incubation with or without DnaK. Clearly, large quantities of amyloid fibrils are observed in the sample containing IAPP alone (Figure 1E). However, aggregation observed in presence of DnaK is minimal (Figure 1F). Thus, Hsp70 appears to inhibit both fibrillar and non-fibrillar growth of IAPP.

Inhibition of the aggregation of IAPP in the absence of ATP indicates that inhibitory action of Hsp70 is not mediated by the catalytic cycle of ‘foldase’ activity rather it is mediated possibly by the ‘holdase’ activity (25). However, interactions of Hsp70 with monomeric IAPP can’t explain the strong effects observed at sub-stoichiometric concentrations (21). Therefore, Hsp70 must interact with one or more on-pathway intermediates of IAPP. The intermediates could be the pre-nucleation oligomers, post-nucleation protofilaments or even ‘misfolded’monomers.

### Inhibitory effect of Hsp70 is the strongest at the prenucleation phase

We then examined whether the inhibitory role of Hsp70 is mediated *via* its interactions with the prenucleation oligomers or with the post-nucleation fibrillar species. Figure 2A shows the time course of aggregation of a 20 μM IAPP solution. DnaK (1 μM) is added into this solution in the beginning (*i.e.*, at t = 0), in the lag phase (at t = 3 h) and in the growth phase (at t = 5 h). The inhibitory effect of DnaK is clearly the highest if it is added at t = 0. The effect is progressively less if added later. Addition of the chaperone in the ‘post-nucleation’ phase slows down the growth of the fibrils but it doesn’t stop it. Figure 2B shows that similar observations are made using Hsc70 as well. Taken together these results indicate that primary mechanism of the inhibitory action of Hsp70 possibly involves prevention of the primary nucleation of IAPP than prevention of the growth of the fibrils.

**Figure 2:**
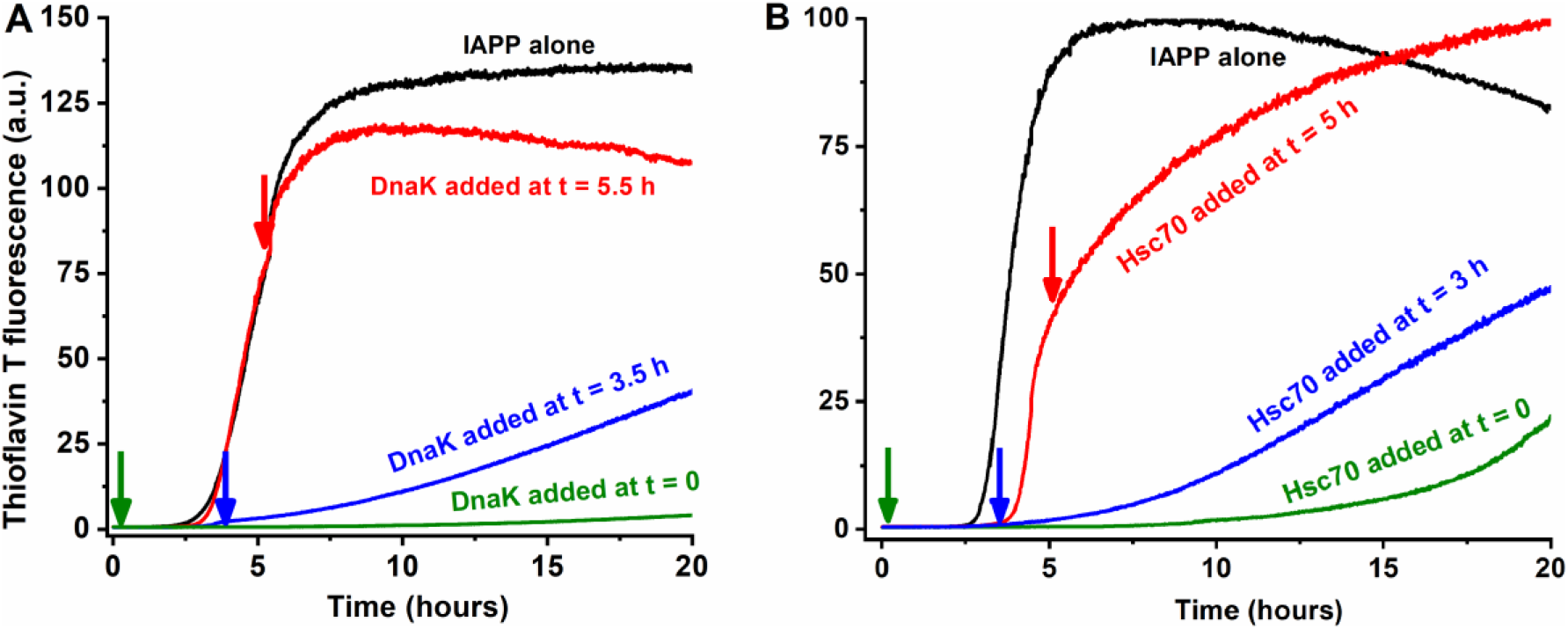
Effect of Hsp70 on different stages of IAPP aggregation. 1 μM DnaK (A) or Hsc70 (B) is added to a 20 μM IAPP solution at different stages of aggregation, *viz*, in the beginning (t=0), ‘lag’ phase (t=3h) and in the growth phase (t=5h). Clearly, the effects of Hsp70 is highest if added early. Solutions are prepared in PBS buffer containing 1 mM EDTA and 1 mM sodium azide at pH 7.4 and incubated at 37 °C with continuous stirring.

### Binding of DnaK with the prefibrillar species of IAPP monitored using Intermolecular Forster Resonance Energy Transfer (FRET)

To monitor intermolecular FRET, we have used EDANS-DnaK and TMR-IAPP. Figure 3A shows that the kinetics of intermolecular FRET exhibit two distinct phases, a fast phase (t < mixing time of the samples, *i.e.*, t < 5 seconds) and a slow phase (t > 1 h). The fast phase most likely indicates binding with the preformed oligomers and/or with the monomers of IAPP. The extremely slow kinetics of the second phase is consistent with binding of DnaK with the oligomers of IAPP that form slowly with time. Therefore, formation of the oligomers is likely the rate limiting in this phase. This is further supported by the observation that the intermolecular FRET reaches completion faster if the concentration of EDANS-DnaK used is lesser, as it can be ‘consumed’ by relatively lower concentration of the oligomers.

**Figure 3:**
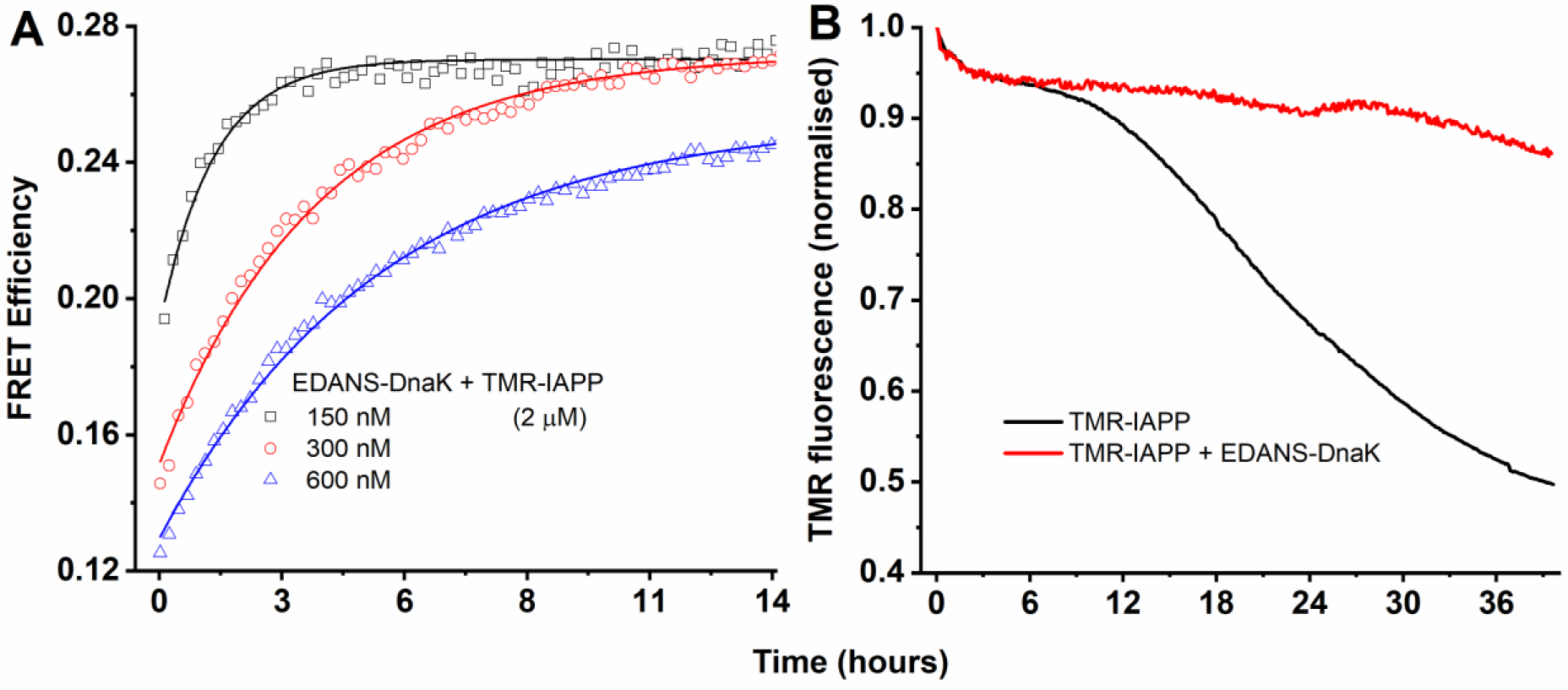
Kinetics of interaction between EDANS-DnaK and TMR-IAPP. A) Kinetics of intermolecular FRET between EDANS-DnaK (150 to 600 nM) and TMR-IAPP (2 μM) monitored using donor (EDANS) fluorescence (λ_ex_ = 350 nm, λ_em_= 480 nm). B) Kinetics of aggregation of TMR-IAPP monitored by fluorescence of TMR in presence and absence of EDANS-DnaK (λ_ex_ = 520 nm, λ_em_= 600 nm). Comparison of the time scales between A and B indicate that the interaction occurs in the early phase of aggregation. The solutions are prepared in PBS buffer containing 1mM EDTA and 1 mM sodium azide at pH 7.4 and incubated at 37 °C with continuous stirring.

To examine the time evolution of the aggregation status of TMR-IAPP, we monitored the fluorescence of TMR under the same experimental conditions following the assay developed by Garai and Frieden (32). The kinetics of fluorescence of TMR-IAPP exhibit in three distinct phases (Figure 3B). These different phases are interpreted as the oligomerization phase (t < 2 h), the ‘lag’ phase (t ~ 10 h) and the growth phase (32). In presence of DnaK, the ‘lag’ phase is extended beyond 30 h. The data presented in Figure 3A indicate that the intermolecular FRET reaches saturation in about 10 h. Thus, intermolecular FRET occurs primarily during the oligomerization and the ‘lag’ phase of aggregation. Therefore, binding of DnaK must involve primarily the prefibrillar species of IAPP.

### Ensemble and single molecule measurements of the binding constant of DnaK/IAPP interactions reflect heterogeneity of the complexes

To determine the binding constant of the interactions, we measured fluorescence anisotropy of TMR-IAPP in the presence of unlabeled DnaK. Since IAPP is a small peptide, anisotropy of TMR-IAPP (MW = 4.4 kDa) is expected to increase considerably upon binding to DnaK (MW = 69 kDa). Figure 4A shows that anisotropy of 400 nM TMR-IAPP increases in presence of 0 to 5 μM DnaK in a concentration dependent manner. While the increase in anisotropy of TMR-IAPP indicates binding to DnaK, the increase is quite small (≈ 20%). The equilibrium constant (*Keq*) obtained from fitting the data using a single site binding model is 2.1 ± 0.9 μM, which appear very high considering that even a few nanomolar DnaK is shown to alter aggregation of IAPP strongly (see Figure 1).

**Figure 4:**
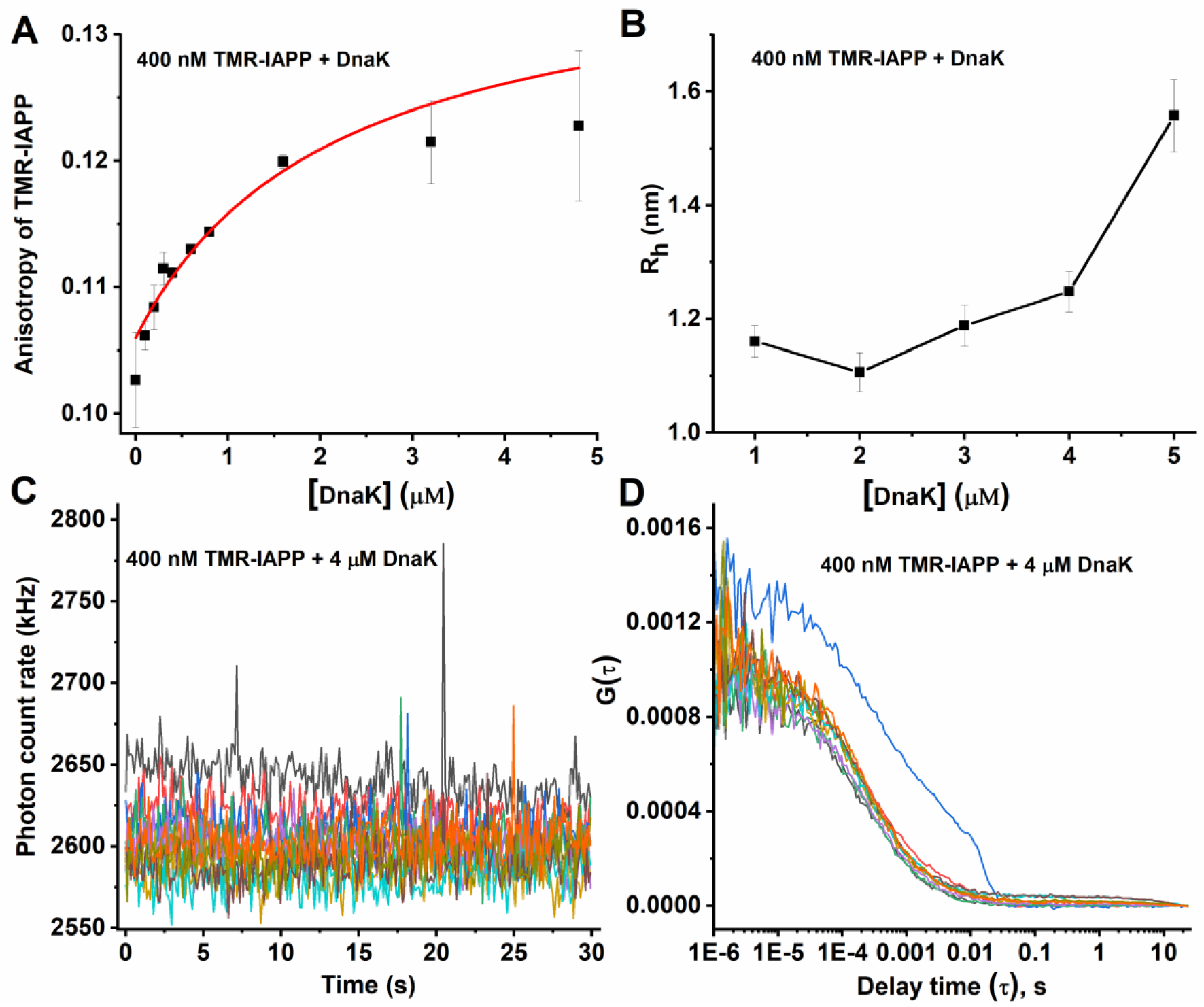
Characterization of the IAPP-DnaK interactions by ensemble and single molecule techniques. A) Ensemble fluorescence anisotropy and B) hydrodynamic radii (R_h_) from FCS measurements of 400 nM TMR-IAPP in presence of 0 to 5 μM DnaK. The symbols represent data. The solid line in A is fit using a single site model. Equilibrium constant obtained is 2.1 μM. Heterogeneity of the system is reflected in the fluorescence bursts in the photon count traces (C) and in differences in the autocorrelation data (D) obtained from 10 identical FCS measurements on 400 nM TMR-IAPP + 4 μM DnaK sample. Fluorescence bursts (C) indicate oligomers. The solutions are prepared in PBS buffer containing 1 mM EDTA and 1 mM sodium azide at pH 7.4 and incubated at RT.

To examine the interactions further we use a single molecule technique, *viz*, Fluorescence Correlation Spectroscopy (FCS) to measure the hydrodynamic radius (R_h_) of TMR-IAPP in presence of DnaK. Since R_h_ is proportional to MW^1/3^, it is expected to increase by ~2.6-times if IAPP binds to monomeric DnaK. Figure 4B shows that R_h_ of TMR-IAPP increases in presence of 0 to 5 μM DnaK in a concentration dependent manner. Surprisingly, the shape of the R_h_ versus [DnaK] plot doesn’t conform to that is expected from a single site binding model as observed in case of the anisotropy data presented in Figure 4A. Furthermore, the increase of R_h_ even in presence of 5 μM DnaK is only about 30%, which is significantly smaller than the expected 160% increase if all the monomeric IAPP are bound to the DnaK. Therefore, most of the IAPP in the solution must be free. Thus, it is possible that DnaK binds primarily to the oligomers of IAPP. Small increase of anisotropy or R_h_ is indicative of the presence of a relatively small population of the oligomers of IAPP. To examine presence of the oligomers, we compared the photon count traces and the autocorrelation traces. Figure 4C and D show typical examples of photon count and autocorrelation traces. Clearly, a significant number of photon count traces exhibit presence of fluorescence bursts indicating presence of oligomers of IAPP. Consequently, large variability is observed among the autocorrelation curves as well. Therefore, FCS data indicate presence of oligomers in the solution. Higher brightness of the oligomeric complexes leads to the heterogeneities in the FCS measurements. We note here that presence of bursts distorts the autocorrelation data (see Figure 4D), hence these data are excluded from the analysis. Thus, nature of the binding curves obtained from the ensemble anisotropy and FCS measurements could differ considerably (see Figure 4A and B).

It may be argued that the above-mentioned heterogeneity is a characteristic of TMR-IAPP. To verify this, we performed FCS measurements of TMR-DnaK in presence of unlabeled IAPP. Many fluorescence bursts are observed in the photon count traces of TMR-DnaK in presence (Figure 5C) but not in absence of IAPP (Figure 5A). The autocorrelation traces obtained in presence of IAPP are also highly heterogeneous as may be expected (Figure 5D). The observed fluorescence bursts are indicative of clustering the TMR-DnaK molecules possibly on the oligomers of IAPP. Quantification of the oligomers in the midst of a vast majority of the free monomers is difficult (34). However, comparison of the photon counting histogram (PCH) of TMR-DnaK in presence and absence of IAPP shows that about 8% of the total photons are associated with the photon bursts (Figure 5E). Hence, about 8% of the 100 nM TMR-DnaK are associated with the complexes giving rise to the bursts. Then we used a photon count threshold of mean + 2 × standard deviation (SD) to count the number of the bursts per minute. We observed about 16 photon bursts per minute. Furthermore, the probability of detecting a photon burst in any time bin (= no. of the bins associated with the bursts/total no. of bins) is found to be about 0.01. In our FCS setup, the average number of molecules (<N>) observed in the confocal volume from 1 nM solution is about 1.1 (35). Hence, the number concentration of the oligomeric complexes must be in the low picomolar range (36). The size of the photon bursts exhibits a broad distribution (Figure 5F). The peak of the distribution corresponds to about 130 kHz. Counts per molecule (CPM) of the monomeric TMR-DnaK obtained from the FCS measurement of the purified TMR-DnaK solution is about 8 kHz (procedure to determine CPM is described in (37). Hence, the average brightness of the complexes are about 16-times higher than monomeric TMR-DnaK. Therefore, oligomers of IAPP interact with high affinity with multiple molecules of DnaK. We note here that the approach outlined above is suitable only for the large bursts, as the small complexes would get masked in the shot noise of the photon counts. Hence, this approach is not suitable for characterization of the small complexes.

**Figure 5:**
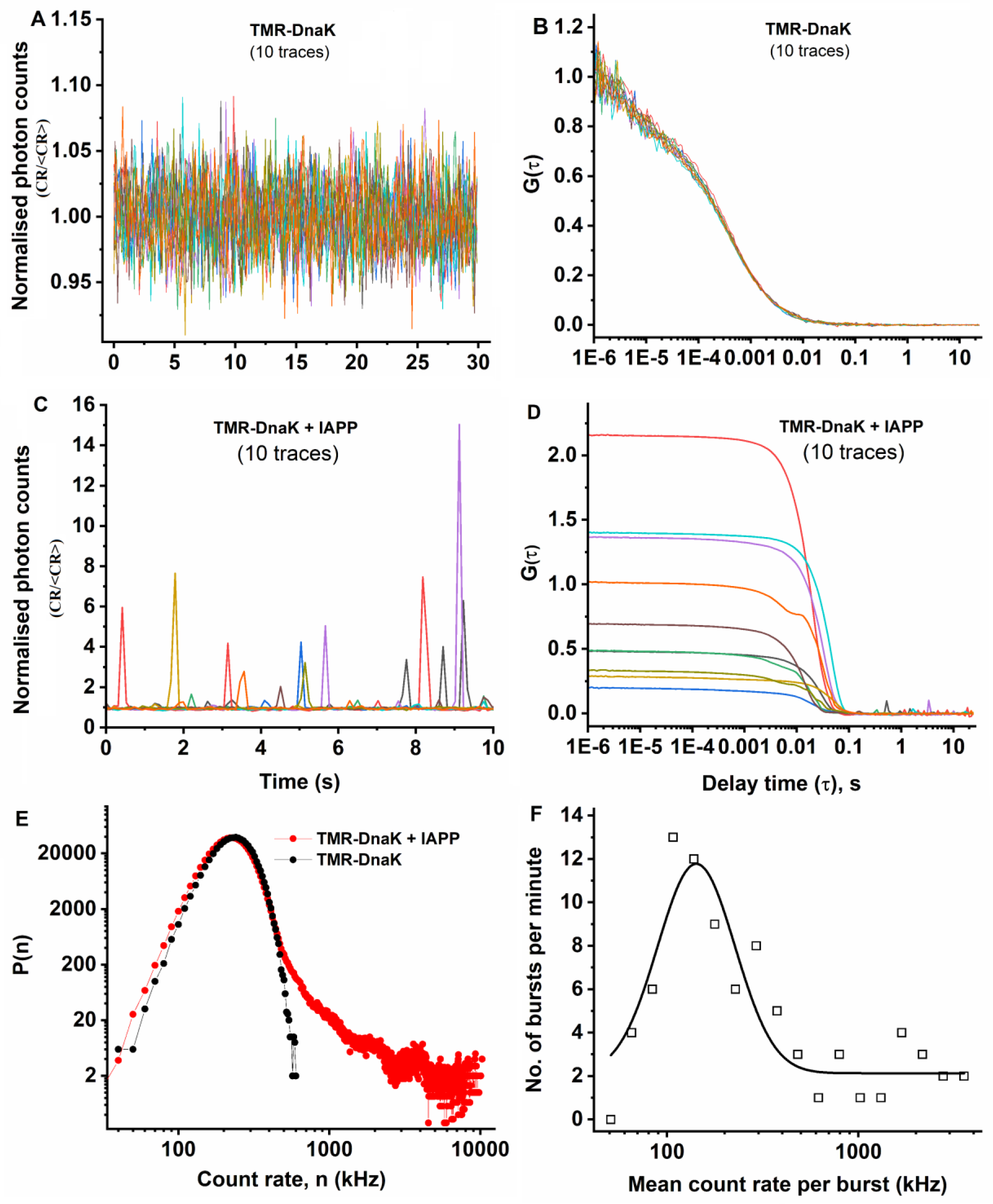
Characterization of the oligomeric IAPP/TMR-DnaK complexes from photon bursts analysis. Photo count traces (A and C) and autocorrelation data (B and D) from 10 identical measurements from 100 nM TMR-DnaK in absence of IAPP (A and B) or in presence of 20 μM IAPP (C and D). Clearly, both photon counts and autocorrelation data are highly heterogeneous in presence of IAPP. E) comparison of photon counting histograms (PCH) indicate about 8% of TMR-DnaK ispartitioned into the bursts. F) histogram of the brightness of the bursts. The solid line is a fit assuming a gaussian distribution. Mean brightness of the bursts (130 kHz) is about 16-times brighter than the monomeric TMR-DnaK. The solutions are prepared in PBS buffer containing 1 mM EDTA and 1 mMsodium azide at pH 7.4 and incubated at RT.

### Characterization of the complexes of DnaK/IAPP by Multiangle Light Scattering (MALS) and FCS following fractionation by SEC and FFF

The FCS measurements presented above indicate that the major problem in characterization of the DnaK/IAPP interaction is heterogeneity of the complexes. Therefore, we use SEC to resolve the various species in the heterogeneous mixture of free IAPP, free DnaK and the IAPP/DnaK complexes. The SEC is connected to a MALS instrument for determination of the absolute molecular weight and the size of the DnaK/IAPP complexes. Figure 6A shows the time course of the concentration of the eluting proteins measured by refractive index (RI) of the solution. It may be seen that the RI (i.e., the concentration) profile shows two closely spaced peaks at 43 and 45 mins. The same figure shows the time course of the molecular weight (MW) obtained from analysis of the MALS data. It may be seen that the MW of the particles eluted corresponding to the peak of the RI is measured to be 70-80 kDa indicating that DnaK exists predominantly as monomers. To verify if the two peaks correspond to complexes of DnaK with different oligomers of IAPP we compared the data obtained from a sample containing DnaK alone. However, the profiles of the RI and the MW of the samples containing DnaK/IAPP and DnaK alone appear to be similar. Then, we performed mass spectrometry measurements on the samples corresponding to the peaks observed in RI to obtain an estimate of the relative concentrations of Dnak and IAPP in the DnaK/IAPP sample. Supplementary figure 1 shows that the peaks contain DnaK. However, no IAPP is detectable in these fractions. This indicates that the population of the DnaK/IAPP complexes must be small. The peaks observed in the RI and the MALS must originate from the primarily free DnaK. The two peaks in RI might indicate slow monomer-dimer equilibrium of DnaK (31). To characterize the DnaK/IAPP complexes specifically we then used FCS. In these experiments we used TMR-IAPP instead of IAPP. Since DnaK used here is not fluorescently labelled the unbound DnaK doesn’t contribute to the signal in FCS. First, SEC is used to fractionate the DnaK/TMR-IAPP complexes from the population of the free TMR-IAPP. Figure 6B shows that the R_h_ of TMR-IAPP/DnaK complex measured using FCS is ~3.3 nm, which is almost same as that of the monomeric DnaK but quite larger than that of the free IAPP (R_h_ ~1.2 nm). Therefore, FCS measurements indicate binding of monomeric DnaK to monomers or small oligomers of TMR-IAPP. Since R_h_ is proportional to (MW)^1/3^, the FCS measurements do not have the resolution to distinguish if these complexes contain monomers or small oligomers of IAPP. However, the concentration of the TMR-IAPP/DnaK complexes detected in FCS is approximately 45 nM (Supplementary Table S1). We note here the concentrations of DnaK and IAPP used in these experiments are several μM. Therefore, concentration of the complexes detected in FCS is very low suggesting that most of the DnaK or the TMR-IAPP in the solution are free.

**Figure 6:**
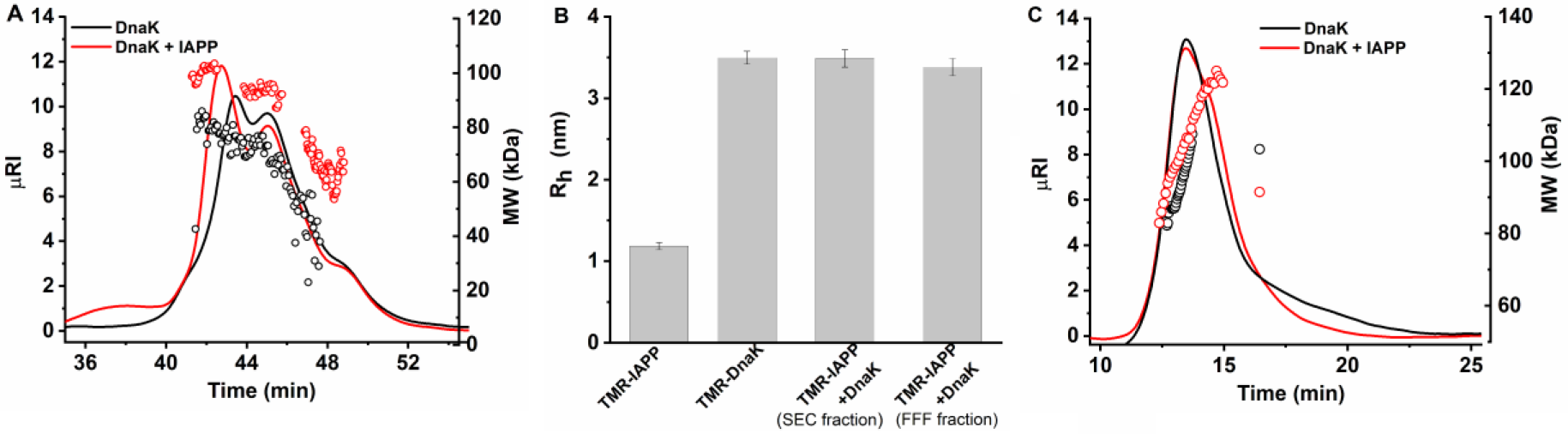
Characterization of IAPP-DnaK complexes following fractionation by Size exclusion chromatography (SEC) or Field flow fractionation (FFF). A) SEC-MALS and C) FFF-MALS of 10 μM DnaK in absence and presence of 20 μM IAPP. The solid lines indicate refractive index (RI), which is a measure of concentration and empty circles represent molecular weight (MW) measured by MALS for DnaK (black) and DnaK+IAPP (Red) samples. B) R_h_ measured using FCS of TMR-IAPP, TMR-DnaK and TMR-IAPP + DnaK following purification using SEC or FFF. Clearly, R_h_ of TMR-IAPP + DnaK is similar to TMR-DnaK indicating binding of TMR-DnaK to small oligomers of IAPP. The solutions are prepared in PBS buffer containing 1 mM EDTA and 1 mM sodium azide at pH 7.4 and incubated at RT.

A well-recognized problem in characterization of non-covalent protein-protein complexes by SEC-MALS is that the complexes may dissociate due to interactions of the proteins with the matrix of the SEC column. To circumvent this problem, we attempt to resolve the various species of the IAPP/DnaK complexes by using Field Flow Fractionation (FFF), which uses flow field to separate the molecules based on hydrodynamic size (38). Figure 6C compares the time courses of the RI and the MW obtained from the mixture of DnaK and IAPP, and the DnaK alone. It may be seen that both the samples show two closely spaced peaks at 12 and 13 mins in RI. The MW of the peaks are found within 60-80 kDa consistent with the SEC-MALS measurements. Once again, we performed FCS measurements on the TMR-IAPP/DnaK samples ‘purified’ using FFF. Analysis of the FCS data presented in Figure 6B show that R_h_ of the complexes are almost same as the monomeric DnaK. Once again, the concentration of the complexes is very low (~ 15 nM) (Supplementary Table S1). Therefore, FFF-MALS data are consistent with the SEC-MALS data. The FCS autocorrelation data and the fits are shown in supplementary figure S2. Taken together, the FCS data indicate that the R_h_ of the complexes are same as the complex of monomeric DnaK binding to monomers or small oligomers of IAPP. Since, the binding of DnaK involves a small fraction of the population of IAPP, we think that DnaK binds to the small oligomers of IAPP.

## Discussion

Amyloid aggregation of proteins is involved in the pathology of multiple human diseases such as Alzheimer’s, Parkinson’s disease and T2DM. Extensive studies over the past three decades indicate that protein monomers first assemble into soluble low molecular weight oligomers, the oligomers then act as sites for nucleation to form small fibrillar aggregates, which grow to larger fibrils. To counteract the numerous pathways that lead to aberrant aggregation of proteins, cells utilise molecular chaperones as a defence mechanism (10,12,39,40). Hsp70 chaperone machinery is the central component of the chaperone systems providing protection against aberrant aggregation of cellular proteins under stress conditions. The commonly depicted mechanism of action is that Hsp70 machinery, which involve the cochaperones such as Hsp40 and the nucleotide exchange factors (NEF), bind to the exposed hydrophobic patches of the fully unfolded polypeptide chains or ‘misfolded’ proteins, and then release it in the correctly ‘folded’ form using an ATP dependent catalytic cycle (41). *In vitro*, Hsp70/Hsp40/NEF machinery has been shown to inhibit aggregation and promote disaggregation of the aggregates of amyloid-β, α-synuclein and polyglutamine polypeptides in presence of ATP (21,25). *In vivo*, over expression or exogeneous addition of Hsp70 has been shown to inhibit amyloid aggregation and rescue cytotoxicity of the amyloids in cell cultures and model organisms (16,20,42).

While the ‘foldase’ activity of Hsp70 is mediated by the ATP dependent catalytic cycle several recent *in vitro* studies have shown that Hsp70 can inhibit amyloid aggregation also in absence of ATP suggesting that the catalytic cycle is not required for the inhibitory action of Hsp70 (12,25). Therefore, Hsp70 can act as a ‘holdase’ in absence of the cochaperones and ATP. This is not surprising as binding of Hsp70 to the polypeptide might delay its aggregation by blocking the sites of growth. However, the extent of the delay would depend on the molecular mechanism of the interactions. For example, if Hsp70 binds predominantly to the monomers then in absence of ATP driven catalytic activity stoichiometric amount of Hsp70 is required to observe significant effects. Our results presented in Figure 1–3 show that Hsp70 can exert strong effects even at substoichiometric concentrations indicating that it must interact with the intermediates/oligomers rather than the monomers of IAPP. This is further supported by the intermolecular FRET measurements which show that DnaK interacts with species of IAPP that form in a time dependent manner (see Figure 3).

However, very little is known so far about the biophysical properties of the DnaK/IAPP complexes. The fluorescence bursts of the TMR-DnaK formed in presence of IAPP suggest that the oligomers of IAPP bind to multiple molecules of DnaK. We find that the concentration of the bursts, i.e., concentration of the complexes is in the picomolar range. Size of the oligomeric complexes could not be determined, however, average brightness of the bursts suggests about 10-100 molecules of DnaK per oligomers. An inherent limitation of the burst analysis in presence of a high background fluorescence, which arises from the large population of the free TMR-DnaK is that the small oligomers cannot be detected or characterized. Therefore, SEC and FFF are used to separate the large oligomeric complexes. We find that the large oligomeric complexes are eliminated by SEC or FFF. While MALS cannot distinguish DnaK/IAPP complexes from free DnaK, FCS can provide information on the size and concentration of the complexes. FCS and MALS measurements on the major fractions indicate that the DnaK/IAPP complexes are small ~90 kDa. Thus, these complexes most likely contain one molecule of DnaK and one or a few molecules of IAPP. However, the concentration of these complexes is very low ~15-45 nM. Since the concentration of the complexes are very low we think that the interaction involves a rare species, i.e., the oligomers of IAPP. Therefore, majority of the DnaK and the IAPP in the mixture must be free. Therefore,our experiments suggest that binding of DnaK with the monomers of IAPP is poor, while affinity for binding to the oligomers is very high.

Our results presented in Figure 2 indicate that the effects of DnaK when added at t = 0 is the strongest. Therefore, DnaK appears to be more effective on the prefibrillar oligomers than the fibrillar forms. Hence, we hypothesize that major role of DnaK is to interact with the oligomers to prevent nucleation rather than preventing growth of the preformed fibrils. The mechanism of the interaction of Hsp70 with substrate proteins has been a matter of intense studies (41,43–46). Hsp70 is known to be highly promiscuous due to its interactions with multitude of substrates with diverse polypeptide sequences (33). However, most of our knowledge about how Hsp70 binds to its substrates is based on the studies using model peptides from experiments using peptide libraries and phage display (47,48). The octapeptide substrate for Hsp70 generally contains a six-residue hydrophobic middle patch flanked by positively charged residues at both ends. Many polypeptide sequences contain such patches. Therefore, Hsp70 binds to fully unfolded polypeptides or the exposed hydrophobic patches in the globular proteins. For example, Hsp70 machinery can refold denatured luciferase *in vitro* (49). Therefore, binding of Hsp70 with globular proteins is regulated by misfolding or unfolding of the protein. However, little is known about how Hsp70 binds to intrinsically disordered proteins (IDP) since the putative binding motifs in IDPs are possibly exposed almost all the time. Therefore, how is the interaction with the IDPs regulated? Molecular chaperone binding site prediction program LIMBO (50) predicts residues 13-19 heptapeptide (ANFLVHS) of IAPP as a probable binding site for DnaK. Therefore, Hsp70 may bind to the patch containing ANFLVHS in the monomeric IAPP. However, our experiments suggest that DnaK binds strongly to the oligomers but weakly to the monomers. Therefore, we think that the high affinity of the interaction with the oligomers arises from multivalent binding. The multivalent binding might involve oligomeric forms of IAPP and the multiple binding sites on Hsp70, similar to what has been proposed earlier for the interaction between apolipoprotein E and the oligomers of amyloid-β (51). While the polypeptide substrates are believed to bind to the canonical substrate binding cleft in the substrate binding domain, the surface of the Hsp70 proteins are capable of interacting with multitude of cochaperone proteins (33). In fact, diversity of the functions of Hsp70 are hypothesised to originate from its ability to interact with diverse array of cochaperones (52). Furthermore, a second binding site that is important for the activity of the Hsp70 has been reported recently (53). The multivalent binding might involve this second binding site and/or other protein-protein interaction sites on the surface of Hsp70. Unlike folded proteins since the IDPs do not possess defined structure, high affinity binding of Hsp70 to the oligomeric but not the monomeric forms of IAPP possibly is required to regulate the interaction ensuring that the cellular functions of the monomeric IAPP are not affected. Furthermore, FCS measurements reported in Figure 5 suggest that oligomers of IAPP bind to multiple molecules of Hsp70. Recruitment of multiple Hsp70 molecules to the protein rhodanase has been hypothesized to promote unfolding of the protein via an ‘entropic pulling’ action (54). Similar mechanism of action may be involved in the disassembly of the oligomers or the fibrils of the amyloid proteins by the Hsp70 machinery (26).

Thus, our data supports that Hsp70 alone can inhibit aggregation of IAPP by the ‘holding’ model (21,25). Binding to the oligomers of IAPP is probably the first step in the catalytic cycle of the chaperone machinery (21,25). Alternatively, Hsp70 can direct the bound complexes to the degradation pathways (55). Several other heat shock proteins such as Hsp90 and Hsp60 also exhibit similar protection against amyloid aggregation (21,56). We find that the Hsp70 binds weakly with the monomers but strongly with affinity in the picomolar range with the oligomeric forms of IAPP. This might be a common feature, which is important for the regulation of the interactions between the constitutively expressed molecular chaperones and the functionally important amyloidogenic IDPs.

## Experimental Procedures

### Preparation of IAPP and Hsp70 chaperones

All the chemicals unless mentioned otherwise were procured from Sigma (USA). Tetramethylrhodamine (TMR)-maleimide and EDANS-maleimide were purchased from Thermo Fischer Scientific (USA).

### Chemical synthesis of IAPP

IAPP was chemically synthesised at 0.1 mmol scale using an automated peptide synthesiser (Aapptec) following a protocol described earlier by Marek et al (57). Crude IAPP was dissolved in 30 mM Acetic acid. The disulfide bond between the cysteines at positions 2 and 7 was induced using hydrogen peroxide. The peptide was stored in the form of lyophilized powder at −20°C. The lyophilized power was dissolved in 10 mM pH 7.4 phosphate buffer containing 4M Guanidine hydrochloride (GdnCl). This solution was purified by size exclusion chromatography using a superdex peptide column (10/300, GE Healthcare, Germany) in 20 mM phosphate pH 7.4 buffer containing 4M GdnCl. Only the monomeric fraction was collected. Finally, a PD10 column (GE Healthcare, Germany) was used to remove the GdnCl. The stock solution was aliquoted into 100 μl vials, flash frozen and stored at −80°C. The mass and purity of the peptide was verified by mass spectrometry (see supplementary figure 3).

### Labeling of IAPP with TMR

Tetramethylrhodamine (TMR) was covalently linked to the N-terminus of IAPP before cleaving the peptide from the resin. Briefly, the N-terminus of IAPP was deprotected using 20% piperidine followed by extensive wash with dimethylformamide (DMF). 5,6-carboxy TMR was then added in 2-fold molar excess and the reaction mixture was kept overnight at RT with stirring. The resin was washed with DMF followed by dichloromethane and methanol to remove free TMR. The peptide was lyophilised and then cleaved from the resin and purified using the procedure mentioned above.

### Expression and purification of *E. coli* DnaK, MTB*-*Hsp70 and Human Hsc70

Plasmids for *E. coli DnaK*, *Mycobacterium tuberculosis Hsp70* and human *Hsc70* are generous gifts from Dr. Shekhar Mande(National Centre for Cell Science, Pune, India), Prof. Lila Gierasch (University of Massachusetts, USA) and Prof. Lewis Kay (University of Toronto, Canada) respectively. Expression and purification of the proteins were performed using the published protocols (46,58,59). BL21 (DE3) *E. coli* cells were transformed with the plasmids described above. Cells were grown in Luria Broth (LB) supplemented with 100 μg/ml ampicillin at 37°C with continuous shaking of 200 rpm. Expression of the proteins was induced with 1 mM IPTG at 0.6 OD of cell density. The cells were harvested by centrifugation after 4 hours of induction. DnaK expressing cells were resuspended in 50 mM Tris pH8.0 and 1 mM EDTA buffer (Buffer A) and lysed using sonication (Branson, USA). The cell lysate was centrifuged at 18000 rpm for 40 minutes using a floor centrifuge (Beckman, USA). The supernatant, which contains DnaK protein was purified first with ion exchange chromatography using 0 to 1 M NaCl in a HiTrap column (HiTrap, GE Healthcare, Germany). Primary fraction of DnaK was eluted in 150 mM NaCl which was further purified using size exclusion chromatography using a superdex 200 column (GE Healthcare, Germany) in PBS, pH 7.4 buffer containing 1 mM EDTA and 1 mM sodium azide. The fraction containing monomeric DnaK was collected, aliquoted into 50 μl vials, flash frozen and stored at −80°C.

Hsc70 and MTB-Hsp70 were purified first by Ni-NTA column chromatography (His-trap, GE Healthcare). The column was washed with 50 mM Tris pH 8, 500 mM NaCl and 20 mM imidazole. The proteins were eluted using 400 mM imidazole. N-terminal His-tag of Hsc70 was removed using TEV protease (1:50 ratio of TEV:Hsc70). The proteins were finally purified using size exclusion chromatography using a Superdex 200 column (10/300 GL, GE Healthcare, Germany) equilibrated with PBS buffer, pH 7.4 containing 1 mM EDTA and 1 mM sodium azide. The pure monomeric fractions of MTB-Hsp70 and Hsc70 were aliquoted in 50 μl vials, flash frozen and stored at −80°C. The purity of the proteins was examined by SDS-PAGE and mass spectrometry (see supplementary figure 4).

### Measurement of kinetics of aggregation of IAPP

A 100 μM stock solution of IAPP was diluted to 20 μM into phosphate buffer saline (PBS) at pH 7.4 containing 4 μM Thioflavin T (ThT) into a round bottom glass test tube (OD = 10 mm) placed inside a temperature-controlled cell holder in the spectrofluorometer (PTI, USA). The solution was stirred continuously using a teflon coated microstirrer (3 mm × 6 mm) bead at approximately 300 rpm at 37 °C. Fluorescence of ThT was monitored continuously using excitation (λ_ex_) at 438 nm and the emission (λ_em_) at 480 nm. Substoichiometric concentrations (125, 250 and 500 nM) of DnaK or MTB-Hsp70 or Hsc70 were added to 20 μM IAPP to examine the effects of the chaperone proteins on the kinetics of aggregation of IAPP. To characterize the effects of Hsp70 on various stages of aggregation, 1μM DnaK (or Hsc70) was added to 20 μM IAPP in the initial (t = 0 h), lag (t = 3 h) and in the growth phase (t = 5 h) of aggregation of IAPP. Aggregation of TMR-IAPP (2 μM) was performed at pH 7.4 in PBS buffer at 37°C with continuous stirring at 300 rpm. The fluorescence of TMR was monitored continuously at λ_em_ = 600 nm with λ_ex_ = 520 nm (32).

### Atomic force microscopy (AFM) imaging

A 10 μl IAPP/DnaK (20 μM/ 0.5 μM) solution was collected from the end of the experiment involving kinetics of aggregation described above. The solution was adsorbed onto a freshly cleaved mica sheet for about 2 minutes. The surface of the mica was then washed gently with Milli-Q water to remove the salt and the loosely bound aggregates. The sheet was then dried in an enclosed chamber for overnight at room temperature. The sample was imaged using an AFM (AFM workshop, CA, USA) in the tapping mode. The resonance frequencies of the cantilever used is within 140–210 kHz and the nominal radius of curvature of the tip used in the measurement is <20 nm. The scanning area is 10 × 10 μm. The scan rate used is 0.5 Hz.

### Kinetics of interaction between DnaK and IAPP by FRET

DnaK was fluorescently labelled at the native cysteine residue at position 6 with EDANS-maleimide using the standard maleimide labeling protocol. EDANS-DnaK was purified using a superdex 200 column (10/300 GL, GE Healthcare, Germany) in PBS buffer at pH 7.4. Different concentrations of EDANS-DnaK (150, 300, 600 nM) were added to TMR-IAPP (2μM) in PBS buffer, pH 7.4. The FRET is monitored continuously using the donor (EDANS) fluorescence using λ_ex_ = 350 nm and λ_em_= 480 nm in the spectrofluorometer (PTI, USA). FRET efficiency was calculated using, FRET efficiency E = 1-F_DA_/F_D_, where F_DA_ is donor emission obtained in presence of the acceptor and F_D_ is donor fluorescence in the absence of acceptor (51). F_D_ was obtained from the fluorescence of EDANS-DnaK monitored in presence of 2 μM unlabeled-IAPP.

### Measurement of DnaK and IAPP interactions using fluorescence anisotropy

Purified 400 nM TMR-IAPP was incubated with different concentrations of DnaK (0 to 5 μM) in PBS buffer (pH 7.4) containing 1 mM EDTA and 1 mM sodium azide. The instrument ‘g’ factor is calculated using free TMR-IAPP. Fluorescence anisotropy of TMR-IAPP was measured in presence of different concentrations of DnaK. The data were fit using a single site binding model using the following equation.

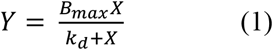

Here, *Y* is the anisotropy, *X* is the concentration of DnaK and *k*_*d*_ is the equilibrium binding constant.

### Fluorescence correlation spectroscopy (FCS) measurements

The FCS experiments were performed using a homebuilt setup described elsewhere (35). We used two excitation lasers, a solid-state laser with λ = 488 nm (Melles Griot, USA) and a He-Ne laser with λ = 543 nm (Newport, USA). The experiments were performed with TMR-IAPP or TMR-DnaK. Hence, only the He-Ne laser was used. For measurement of binding of TMR-IAPP and DnaK, TMR-IAPP (400 nM) solution was incubated with 0 to 5 μM DnaK in PBS buffer, pH 7.4 in presence of 1 mM EDTA and 1 mM NaN3 at RT. The autocorrelation data was analysed using a single component diffusion model. Presence of fluorescence bursts distort the autocorrelation data. Hence, these are removed from the analysis. The diffusion time obtained was used to calculate the hydrodynamic radii (R_h_) (35). Calibration of the FCS observation volume was performed using free rhodamine B (37).

### Burst analysis

Photon count traces obtained from the FCS measurements of 100 nM TMR-DnaK in presence of 20 μM IAPP were recorded with a bin time of 100 μs. First, the photon counting histogram (PCH) is calculated. To obtain an estimate of the mean and the width (SD) of the PCH, the data are fit using a gaussian function. We note here that the appropriate function to fit the PCH data is quite complex. Since our objective is to estimate the mean and the SD only to decide a threshold for detection of the bursts, we have used fitting of the PCH simply with a Gaussian function. Analysis of the fluorescence bursts were performed using approaches similar to those described in (60). The threshold is set at mean + 2 × SD. The number of bursts, total photon counts (*n*) per burst and the temporal size (τ) of the bursts are calculated from the raw photon count data using the free available module ‘peakutils’ in Python. The brightness of each burst is calculated as, ϵ = *n*/τ.

### FFF-MALS and SEC-MALS measurements

The FFF and the MALS instruments are from Postnova Analytics (Germany). The FFF was used to separate the DnaK/IAPP complexes from the free IAPP. The MALS is connected in-line with the FFF for the measurements of the molecular weight (MW) and radius of gyration (R_g_) of the proteins. IAPP (20 μM) was incubated with 10 μM DnaK in PBS pH 7.4 buffer at RT. A 100 μl aliquot of this mixture was injected in the FFF system. We have used a 5 kDa cellulose membrane, 0.20 ml/min focus flow for 6 minutes, 0.25 ml/min forward flow and 3 ml/min cross flow in the FFF. The dn/dc measurements were performed using a refractive index (RI) detector (Postnova Analytics). The SEC-MALS measurements were performed in the same setup. In this case instead of using the FFF membrane assembly a Superdex 200 column was used for fractionation of the DnaK/IAPP complexes. The flow rate used in SEC is 0.4ml/min.

## Acknowledgements

We acknowledge funding from Department of Atomic Energy, India. This work is partly supported by early career grant received by KG from Science and Engineering Research Board, India.

## Conflict of Interest

The authors declare no conflict of interest.

## Author contributions

KG conceptualized research. NC, BS, DS, SC and MM performed research. KG and BS performed analysis. KG and NC wrote the paper.

## Supporting information

### Mass spectrometry of the TMR-IAPP/DnaK solution following fractionation by Size exclusion Chromatography

TMR-IAPP/DnaK complexes were fractionated using size exclusion chromatography (SEC), which is coupled to the multiangle light scattering (MALS) setup. The fraction exhibiting the highest peak in the refractive index (RI) detector was collected for mass spectrometry measurements. Mass spectrometry measurements were performed using a 6545 Q-TOF ESI-MS (Agilent technologies). The spectrometer is coupled to a HPLC for desalting of the injected sample. The m/z spectra were deconvoluted with the MassHunter software (Agilent, UK) using the resolved isotope deconvolution method for measuring the mass of peptides (MW < 10 kDa), and the maximum entropy deconvolution method for measuring the mass of proteins (MW > 10kDa).

**Supplementary Figure S1:**
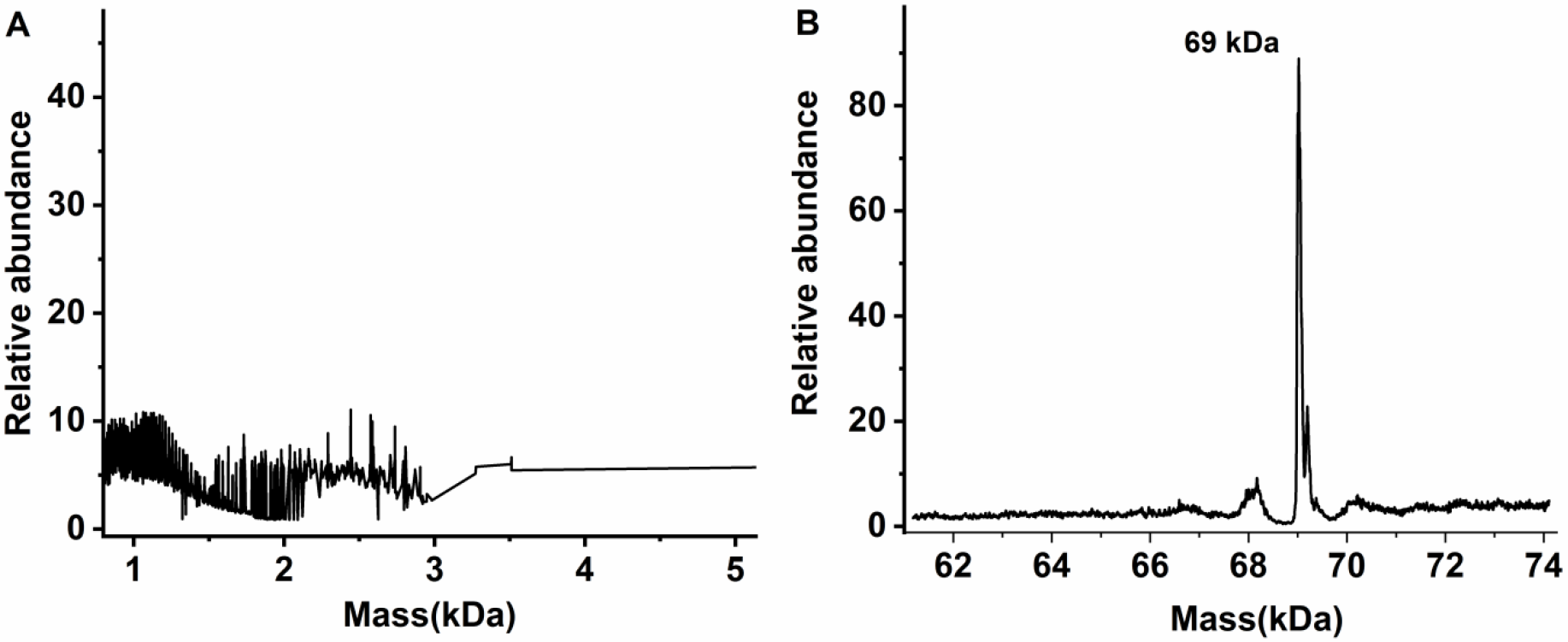
Mass spectrometry of the TMR-IAPP/DnaK sample following fractionation by SEC. A) Deconvolution of the m/z spectra of IAPP was performed within a mass range of 0 to 5000 Da using the resolved isotope method. Deconvolution within this mass range did not detect any IAPP, indicating that concentration of IAPP in the SEC fraction must be very low. B) Detection of DnaK by deconvolution of m/z spectrum within a mass range of 60 to 80 kDa. Presence of DnaK (69 kDa) is quite clear. Hence, the SEC purified fraction contains primarily free DnaK and possibly a very low concentration of the complexes.

### Autocorrelation data G(τ) obtained from the FCS measurements of SEC or FFF purified TMR-IAPP/DnaK

FCS measurements were performed on free rhodamine B, TMR-IAPP, TMR-DnaK and SEC or FFF purified fractions of TMR-IAPP/DnaK solution. Free rhodamine B (50 nM) was used as a control to characterize the FCS observation volume. TMR-IAPP (50 nM) and TMR-DnaK (50 nM) were used to estimate the hydrodynamic radii (R_h_) of these proteins in the unbound monomeric forms. The major fractions of the SEC or FFF purified TMR-IAPP/DnaK solution was used directly for FCS measurements. The FCS data were analysed using a single diffusion and single relaxation model (1) (see supplementary eq. S1). The relaxation component was used to take care of the contributions from conformational fluctuations and/or the triplet state dynamics. Concentrations of the purified samples were determined from the G(0) of the autocorrelation data..

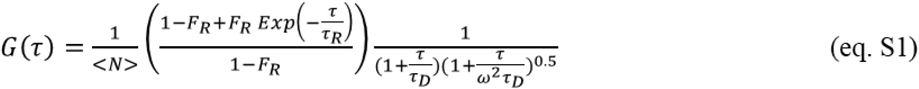

Here <N> is average number of molecules in the FCS observation volume. F_R_ and τ_R_ are the relative amplitude and the characteristic time of the relaxation component respectively. The τ_D_ is the diffusion time of the molecule and ω is the axial ratio of the FCS observation volume. It may noted here that <N>^−1^ is G(0)×(1-F_R_). Hence, in absence of the relaxation component <N>^−1^ is simply equal to G(0). Hydrodynamic radius (R_h_) of the proteins are estimated from the measured diffusion times using the following relationship (see eq. S2). The R_h_ of rhodamine B used is 0.57 nm (2).

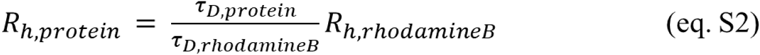

**Supplementary Figure S2:**
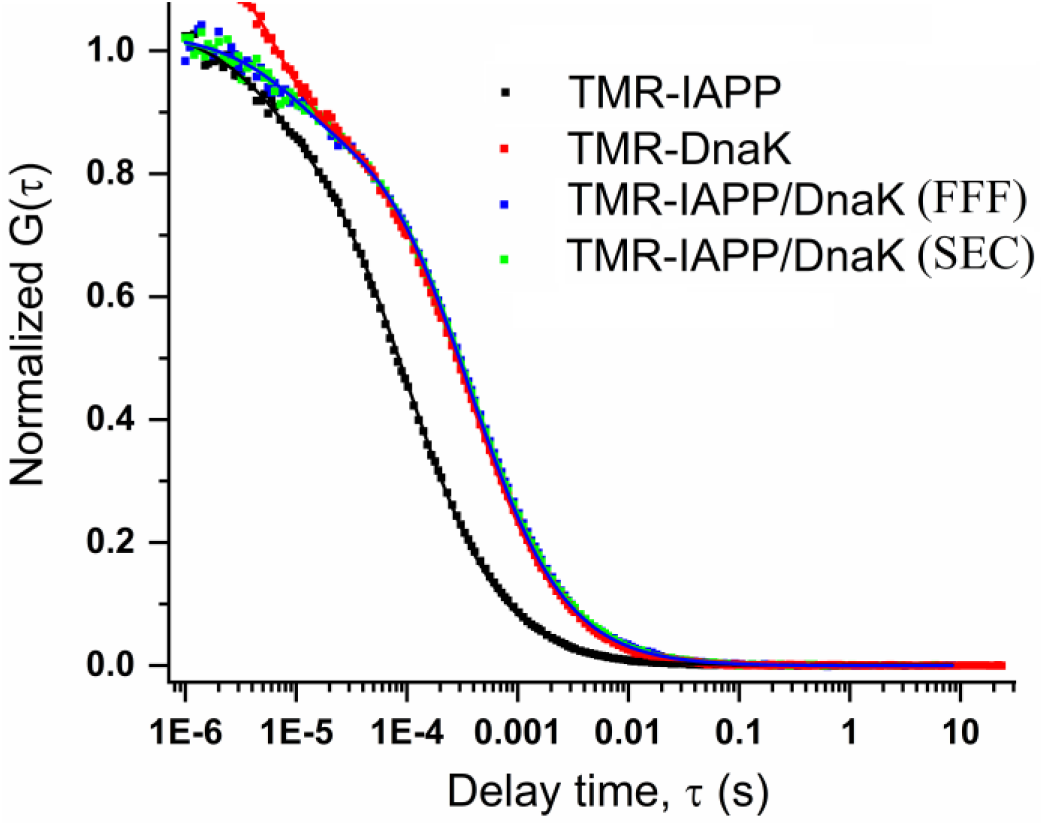
Normalized autocorrelation data G(τ) obtained from FCS measurements of SEC or FFF purified IAPP/DnaK. TMR-IAPP (black), TMR-DnaK (red), SEC purified fraction (green) or FFF purified fraction (blue) of the TMR-IAPP/DnaK sample in PBS, pH 7.4 buffer at RT. For SEC or FFF purified samples, the fractions exhibiting the highest RI were taken for the FCS measurements. The symbols represent data and the solid lines were fit using a single diffusion and single relaxation model (see supplementary eq. S1). The diffusion time (τ_D_) was used for estimation of the hydrodynamic radius of the molecules (see supplementary eq. S2).

**Supplementary Table S1:**
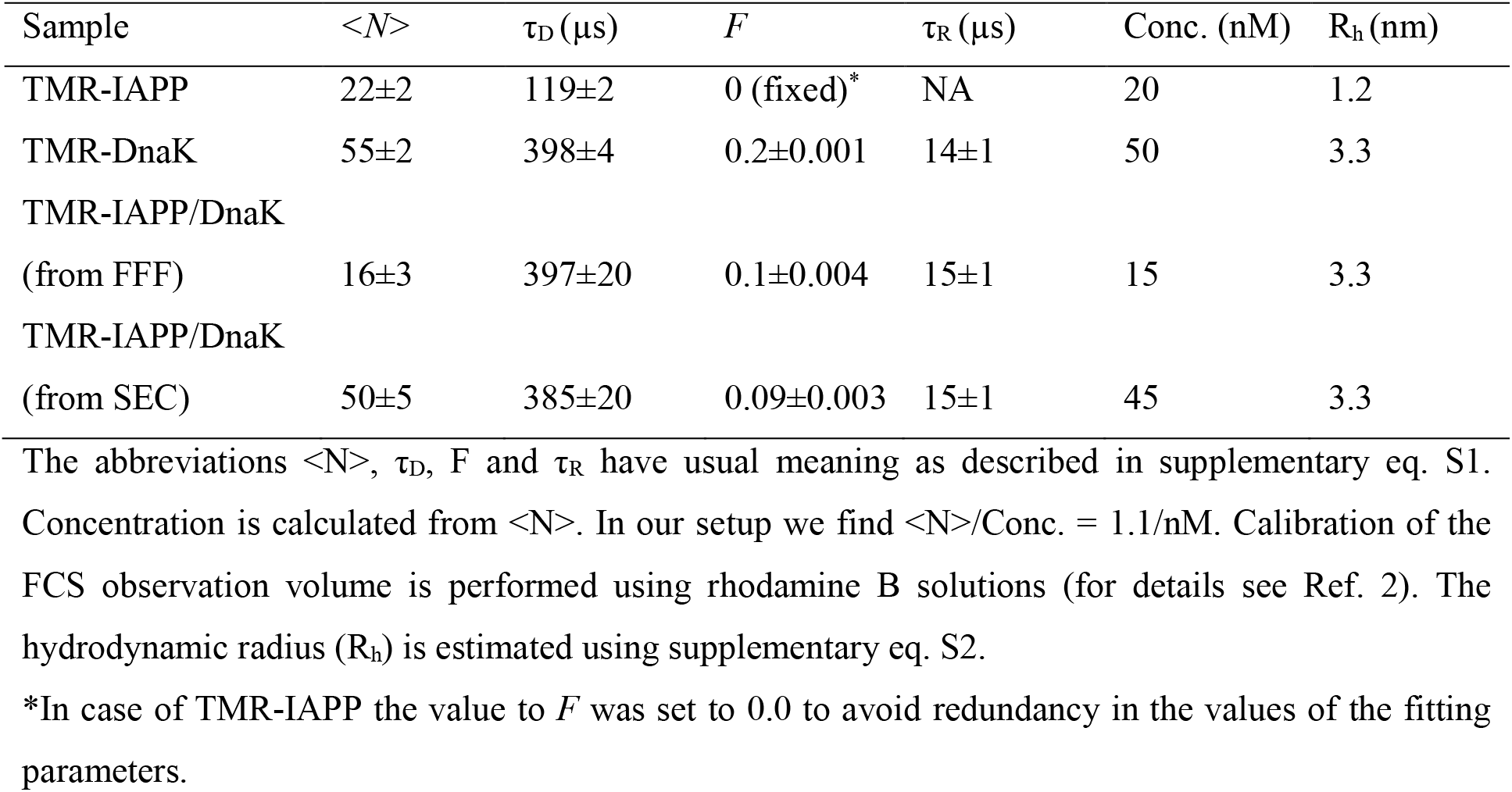
Summary of the parameters obtained from fitting of the FCS data.

### Mass spectrometry of chemically synthesized purified IAPP

A 100 μl of IAPP sample was injected into the HPLC fitted with reverse phase column (Advance Biopeptide map column, Agilent), which is connected inline with an ESI-MS mass spectrometer (Agilent technologies, UK). The m/z spectra were deconvoluted using the inbuilt program in the resolved isotope mode and mass of 3904.9 Da was noted for IAPP which is similar to that of the theoretical MW of IAPP containing disulphide bridge between 2 and 7 cysteine.

**Supplementary Figure S3:**
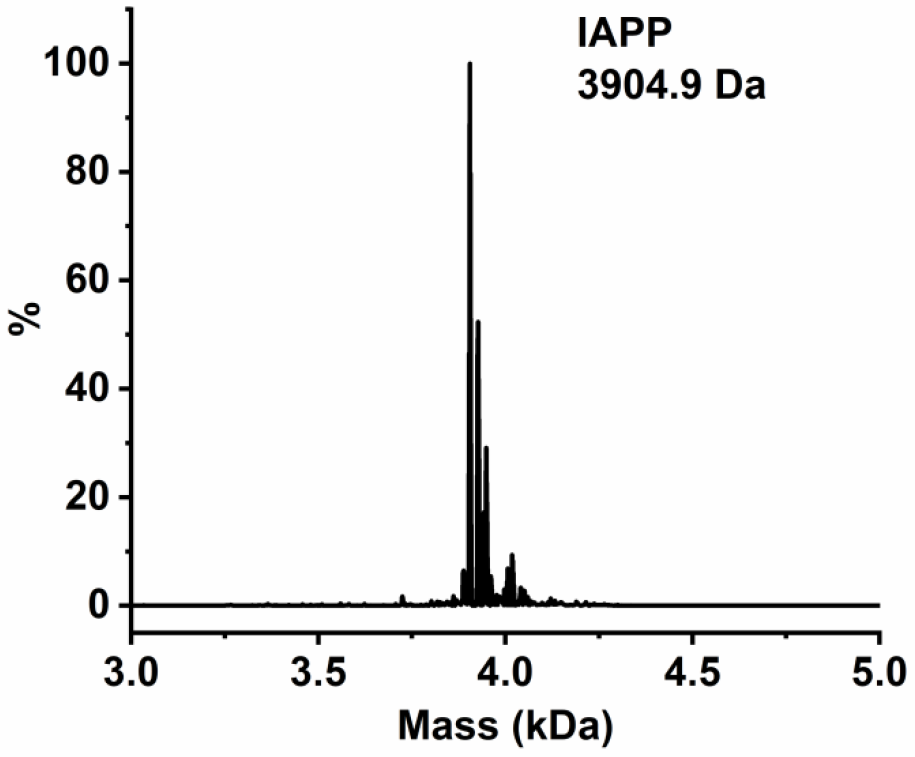
Mass spectrometry of chemically synthesized purified IAPP. Molecular weight (MW) and the purity of the chemically synthesized IAPP were confirmed by mass spectrometry. The MW of IAPP was measured to be 3904.9 Da, which is same as the theoretical MW of IAPP containing disulphide bridge between cysteines placed at position 2 and 7. MW of the peaks placed on the right side of the main peak are 3925.2 and 3949.2 Da. These peaks are corresponding to the sodium adducts of IAPP.

**Supplementary Figure S4:**
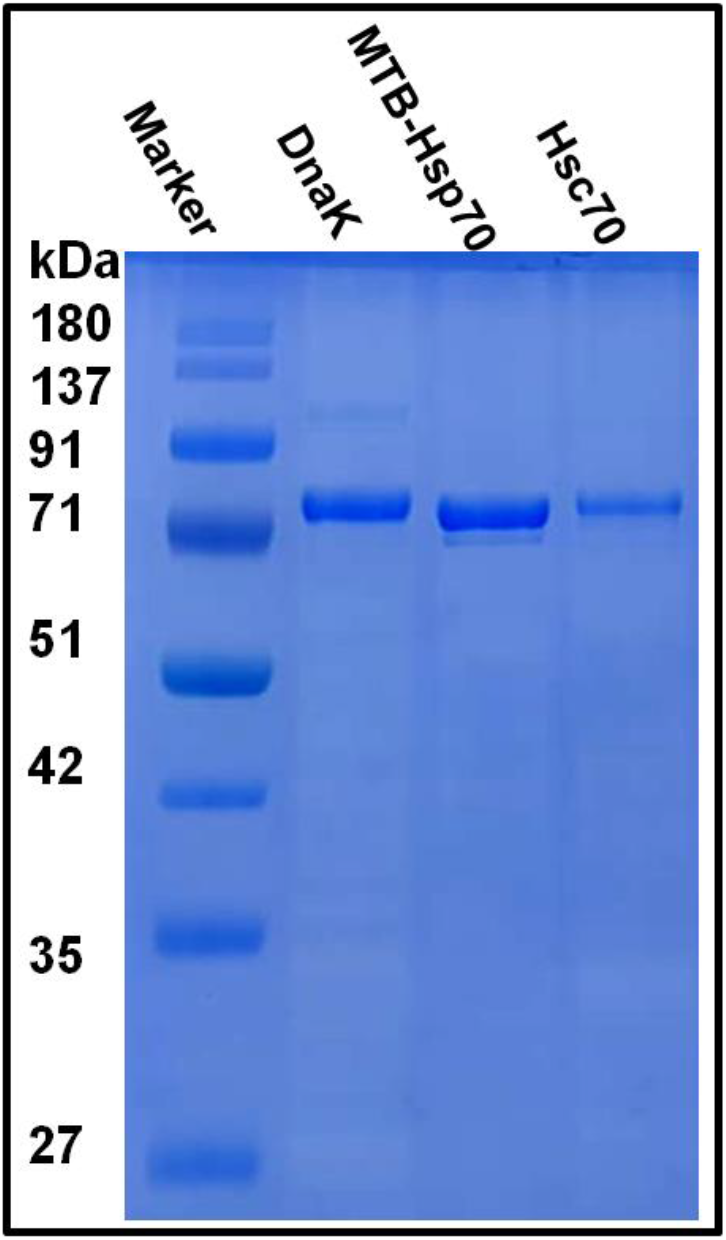
SDS-PAGE of purified *E. coli* DnaK, MTB-Hsp70 and human Hsc70. A single dominant band indicates that the proteins are considerably pure.

